# Role of the lipolysis stimulated lipoprotein receptor LSR in the regulation of macrophage lipid homeostasis and metabolic activity

**DOI:** 10.64898/2026.10.16.759558

**Authors:** Gaëtan Vanotti, Noah Kleinberg, Huguette Louis, Fatiha Maskali, Magnus Bäck, Nathalie Mercier, Frances T. Yen

## Abstract

In atherosclerosis, monocyte-derived macrophages in the subendothelial space of the arterial wall, become lipid-laden and pro-inflammatory. Macrophages depend on lipoprotein receptors to regulate lipid influx, thereby shaping macrophage metabolic state and potentially dictating their phenotype. The lipolysis stimulated lipoprotein receptor, LSR, contributes to lipid homeostasis and was detected by immunofluorescence in aortic atherosclerotic plaques from ApoE-/- mice and by immunoblot in macrophages. The aim was to determine LSR’s role in macrophage lipid handling by studying LSR expression and activity before (M0) or after polarization of Raw 264.7 cells using LPS/IFNγ (M1) or IL-4/IL-13 (M2), respectively. LSR mRNA (0.48-fold, p < 0.001), protein levels (0.28-fold, p < 0.001) were significantly lower in M1 (iNOS+/MCP1+) as compared to M2 (CD206+/Arg1+) macrophages. LSR activity, measured as oleate-induced uptake of DiI-labelled LDL, was significantly lower in M1 as compared to M2 macrophages. siRNA-mediated LSR knockdown (87±4%, p < 0.01) in M0 cells led to significantly decreased expression of genes involved in lipid influx (LDL-R, LRP1) and transport (ABCA1), increased basal glycolysis and decreased glycolytic reserve, but without changes in inflammatory markers. Raw 264.7 macrophages transformation into foam cells by 24h incubation with oxidized LDL also led to reduced LSR activity and expression. We propose that the lower LSR expression and activity observed in M1 and foam cell macrophages contribute to reduced lipid influx and increased glycolytic state that is associated with pro-inflammatory M1 phenotype. This provides new insight towards understanding how these immune cells depend on lipoprotein receptors regulating their metabolic uptake.

## Introduction

Atherosclerosis in its early stage involves endothelial dysfunction, infiltration of monocytes and lipoproteins such as LDL from the circulation which become trapped in the subendothelial space of the arterial wall [1]. Lipids accumulate in the vessel wall leading to formation of fatty lesions, while the local tissue environment sends out pro-inflammatory signals recruiting immune cells [1]. Monocyte-derived macrophages play a key role in the initial stage of this chronic inflammatory disease, becoming lipid-laden and transforming into foam cells to become the core of the atherosclerotic plaque. Macrophages exhibit high plasticity, allowing them shift to their phenotype in response to microenvironmental stimuli. Although heterogeneous, macrophage phenotypes are broadly classified into pro-inflammatory M1 macrophages—induced by lipopolysaccharide (LPS) and interferon-gamma (IFN-γ) and associated with plaque progression—and anti-inflammatory M2 macrophages, which are induced by cytokines such as IL-4 and IL-13 to promote tissue repair and reduce inflammation [2].

By virtue of their ability to adapt to the local tissue environment, the macrophage plays a key role in atherosclerotic plaque formation. How these immune cells adapt their function and phenotype in response to their surrounding environment may be linked to their metabolic status. Lipid homeostasis or the regulation of the balance between lipid synthesis, uptake, storage and catabolism is essential for cell metabolism and function. Indeed, M1 show increased glycolysis and lipid storage, while M2 macrophages metabolism shifts towards increased oxidative phosphorylation in part due to higher fatty acid oxidation and uptake [2]. To ensure proper lipid handling and storage, macrophages rely on lipoprotein receptors, such as the LDL-receptor (LDL-R) family [3,4] that act as “gatekeepers” in regulating the internalization of lipoproteins, and thus the influx of lipids. Foam cell formation has been mainly attributed to the scavenger receptor (CD36 and SR-A1)-mediated uptake of oxidized lipoproteins [5]. Because these receptors are not downregulated by high intracellular cholesterol levels [6], the uncontrolled cellular uptake of oxidized LDL (oxLDL) causes a shift towards lipid storage of lipids in the form of cholesteryl esters, thereby contributing to foam cell formation.

We identified the lipolysis stimulated lipoprotein receptor (LSR), which is involved in the regulation of lipid homeostasis [7]. This receptor is a multimeric protein complex on the plasma membrane composed of 3 subunits α, α’ and β derived by alternative splicing from the same gene [8,9]. It is unique to other lipoprotein receptors in that it requires fatty acids to bind and internalize ApoB (LDL) or ApoE (VLDL, chylomicrons)-containing lipoproteins [7,10]. Transient transfection studies showed that at least the α and β are required for delivery of the lipoproteins to the lysosome for degradation following LSR-mediated endocytosis [8]. Hepatic LSR has been shown to contribute to the clearance of both VLDL and LDL, as shown by the dyslipidemia developing in heterozygote *lsr+/-* mice [10] or following siRNA-mediated hepatic LSR inactivation [11]. In addition, *lsr+/-* mice placed on a high-fat high-cholesterol diet show increased lipid accumulation in aortas suggesting a potential role of LSR in lipid accumulation and plaque development at the level of the vessel wall [10]. The goal of this *in vitro* study was therefore to determine LSR’s role in macrophage lipid handling before (M0) or after polarization of Raw 264.7 macrophage cell line into M1 or M2.

## Materials and Methods

### Cells

Mouse Raw 264.7 macrophage cells (ATCC TIB-71) were cultivated in high-glucose DMEM (Merck KGaA, Darmstadt, Germany) containing 10% heat-inactivated fetal calf serum (Merck KGaA), 1 g/L L-glutamine (Merck KGaA) and 1 g/L penicillin/streptomycin (Merck KGaA), and used at low passages (max 18)[12]. The cells were maintained in culture at 37°C in a humid atmosphere with 5% CO₂. Aortic vascular smooth muscle cells (VSMC) and endothelial (EC) cells were graciously provided by Dr Véronique Regnault.

### Animals

Nine-week-old C57BL6/J and ApoE-/- mice (Charles River, Miserey, France) were placed on a chow or high-fat diet (SAFE® WESTERN 1635; 0.2% Cholesterol, 21% fat from dairy butter; Aston Pharma, London, UK) for 16 weeks. Mice were housed in certified animal facilities under standard conditions with access *ad libitum* to food and water. All procedures were approved by the local ethics committee and the Ministry of National Education, Higher Education, and Research and Innovation in France (Ethical permit n° APAFIS-2024040922135882-v4). At the end of the study, aortas were removed, and cryosections were immediately prepared and fixed for immunofluorescence studies.

### Polarization of macrophages

Raw 264.7 cells (M0) were seeded at 150,000 cells/well in 6-well plates or on coverslips and incubated for 24h at 37°C in a humid atmosphere containing 5% CO2, then incubated 72h at 37°C with lipopolysaccharide (LPS, 100 ng/mL) and interferon γ (IFNγ, 20 ng/mL) for the M1 phenotype or with interleukin 4 (IL-4, 20 ng/mL) and interleukin 13 (IL-13, 20 ng/mL) for the M2 phenotype [13].

### Treatment of cells with siRNA directed against lsr

Raw 264.7 (M0) cells were seeded at 50,000 cells per well in 6-well plates. After 24 h incubation at 37°C, the cells were then treated with control siRNA or an *lsr*-targeted siRNA (Ambion, Thermofisher)[14] using Interferin (Polyplus, Illkirch-Graffenstaden, France) as transfection agent, according to manufacturer’s instructions, followed by incubation for 3 days at 37°C.

### Metabolic studies (Seahorse®)

After siRNA treatment, Raw 264.7 cells were harvested and then reseeded into 8-well culture plates for the Seahorse® XFp. After 24h incubation at 37°C, medium was changed in order to perform the glycolytic activity, glycolytic stress and ATP production assays according to the manufacturer’s instructions. Three independent runs were performed in triplicate; data was analyzed using Wave Seahorse® software. Data were normalized to corresponding controls for each independent run.

### Preparation, labeling and oxidation of LDL

Human LDL (1.025 < d < 1.055) was prepared by sequential ultracentrifugation from plasma obtained at the local blood bank, as described previously [10], filtered (0.22 µ), and then stored at 4°C under N2(g), and used within 8 days. LDL was labeled with the fluorescent probe DiI [10], and used within one week of its preparation. LDL protein was determined using the modified Lowry assay [15]. LDL in 0.15M NaCl was minimally oxidized by UV exposure (Crosslinker CL-508, Uvitec, Cambridge, 0.5 mW/cm², 254 nm), followed by addition of EDTA (0.24 mM, final concentration) to stop the oxidation. Oxidation was verified by measuring thiobarbituric acid reactive substances (TBAR) levels as described previously [16]. Lipoprotein deficient serum (LPDS, d < 1.25 g/mL) was prepared by ultracentrifugation [14].

### Foam cell formation

Cells in 6- or 24-well plates at 70-80% confluence were washed in PBS and then incubated 4h at 37°C _2_ with medium containing 0.2% BSA and 5 mM Hepes for 4h, followed by incubation for 24h at 37°C with medium containing 10% (v/v) LPDS and 100 µg/mL of LDL or oxidized LDL (LDL was oxidized on the same day of treatment).

### Lipoprotein internalization studies

After polarization or foam cell formation as described above in 24-well plates, cells were rinsed with PBS, and incubated with DiI-LDL in DMEM media containing 0.2% BSA and 5 mM HEPES, pH 7.5 supplemented or not with 0.1 mM oleate (from oleate stock solution in isopropanol). Control cells received isopropanol alone at < 1%. One set of cells received 20-fold excess unlabeled LDL to measure nonspecific binding. After 2h incubation at 37°C in 5% CO_2_ incubator, cells were washed once rapidly, and then two 10 min washes with ice-cold PBS/0.2% BSA, followed by two rapid washes with ice-cold PBS alone. Cells were then lysed by 20 min at room temperature with 0.1N NaOH (200 µL/well). Cell lysates were recovered and fluorescence was measured on a microplate reader (CLARIOstar Plus, BMG Labteck, Ortenberg, Germany). The amount of DiI LDL that was internalized was calculated using a standard curve prepared with the same Dil-LDL preparation. Protein levels of the cell lysates was determined using the Pierce™ BCA Protein Assay Kit (ThermoFisher Scientific) according to the manufacturer’s instructions.

### Immunofluorescence (IF) labeling and analysis

Cells on coverslips after polarization were fixed 5 min at room temperature with 4% paraformaldehyde (PAF), followed by three 5-minute washes with PBS. Permeabilization and saturation was performed by a 10 min incubation with PBS containing 5% (w/v) BSA and 0.2% (v/v) Triton, followed by a 55 min incubation with PBS and 5% BSA. Cells were incubated at 4°C overnight with primary antibodies (**Supplementary Table 1**) in PBS containing 1% (w/v) BSA. Coverslips were washed, incubated with conjugated secondary antibodies in PBS containing 1% BSA for 90 min at room temperature. After washing, coverslips were mounted on slides mounting medium containing DAPI. Images were taken using a Nikon Eclipse CI epifluorescence microscope equipped with a Nikon Intensilight C-HGFI lamp and a Nikon D5-Qi2 camera. The fluorescence images were processed and analyzed using FIJI software (ImageJ, Rasband W.S.).

### Western blot and analysis

Treated cells were recovered in lysis buffer (Complete LysisM, Roche, Basel, Switzerland). After 1h incubation at 4°C, samples were centrifuged for 30 minutes at 10,000 g at 4°C, and the supernatant was recovered for Western blots. Protein content of cell lysates was determined using the Pierce™ BCA Protein Assay Kit (Thermo Scientific). Protein (25 µg/well) and pre-stained MW markers (All blue, BioRAd) were separated on 10% SDS-polyacrylamide gels. Gels were recovered and activated by UV for 45 sec (Fusion FX Imager, Vilber, Marne-la-Vallée, France) to obtain a profile of the total amount of protein on the gel. Proteins were transferred to nitrocellulose membranes (Trans-Blot® TurboTM System, BioRad, 10 min, 25 V, 2.5 A). Transfer of proteins was verified by UV, and then stained with Ponceau Red. Membranes were washed with Tris-buffered saline (TBS) containing 0.1% (v/v) Tween 20 (TBS-T). Membranes were blocked by 1h incubation at room temperature with TBS-T containing 5% milk, and then incubated overnight at 4°C with the primary antibodies. After three washes, membranes were incubated 90 min at room temperature with the corresponding secondary antibodies. After three washes, bands were detected using the Clarity™ kit (170-5060, BioRad, Hercules, USA) according to the manufacturer’s instructions with the Fusion FX imager.

### qPCR

Treated cells were washed with PBS, and then recovered in 350 µl of lysis buffer reagent after 10 min shaking at 4°C. After total RNA extraction (Nucleospin RNA Plus kit, Machery-Nagel, Hoerdt, France), RNA was heated to 65°C for 5 minutes to obtain single-stranded RNA. RNA concentration and sample purity was verified as A260/A280 nm and A260/A230 nm ratios falling between 1.8-2.2 (Nanodrop One, Thermo Fisher Scientific, Waltham, United States). cDNA was prepared from total RNA using the iScript reverse transcription (RT) kit (BioRad, Marnes-la-Coquette, France), and used for teverse transcription under the following conditions: 25°C for 5 min, 46°C for 20 min, and 95°C for 1 min. cDNA diluted 1:5 was added to a SybrGreen master mix (Bio-Rad) and primers (**Supplementary Table 2**; final concentration, 0.3 µM). RT-PCR was performed on the BioRad CFX Connect-Real-Time System (BioRad) using the following conditions: 3 min denaturation at 95°C, 40 cycles of 95°C at 10 sec, hybridization at the appropriate temperature, 30 sec at 72°C for primer extension, annealing at 95°C for 10 sec, and then at 65°C for 5 sec, and inactivation at 95°C for 5 sec. β-actin was used as reference genes. Relative expression of genes was analyzed using the 2^-ΔΔCt^ method.

### Statistical analysis

All statistical tests were performed using GraphPad Prism 11 software. Results are shown as mean ± SD or SEM as indicated. The normality of the distributions was assessed using: Anderson-Darling, D’Agostino-Pearson omnibus, Shapiro-Wilk, and Kolmogorov-Smirnov tests. For internalization assays, comparisons between the groups with and without oleate were performed using Student’s t-test. Comparisons of polarization markers were performed using one-way ANOVAs; p-values < 0.05 were considered statistically significant.

## Results

IF revealed LSR expression in both the intima (endothelial cells, EC) and the media (vascular smooth muscle cells, VSMC) of the murine aortic wall (**Fig. 1A**). Immunofluorescence staining of atherosclerotic plaques found in aorta sections from 25-week-old ApoE-/- mice after 16 weeks on Western diet also revealed LSR localization in atherosclerotic plaques (**Fig 1A, right panels**). Strong intensity of LSR labeling (red) was observed in the plaque that did not co-label with the EC or VSMC.

**Figure 1.**
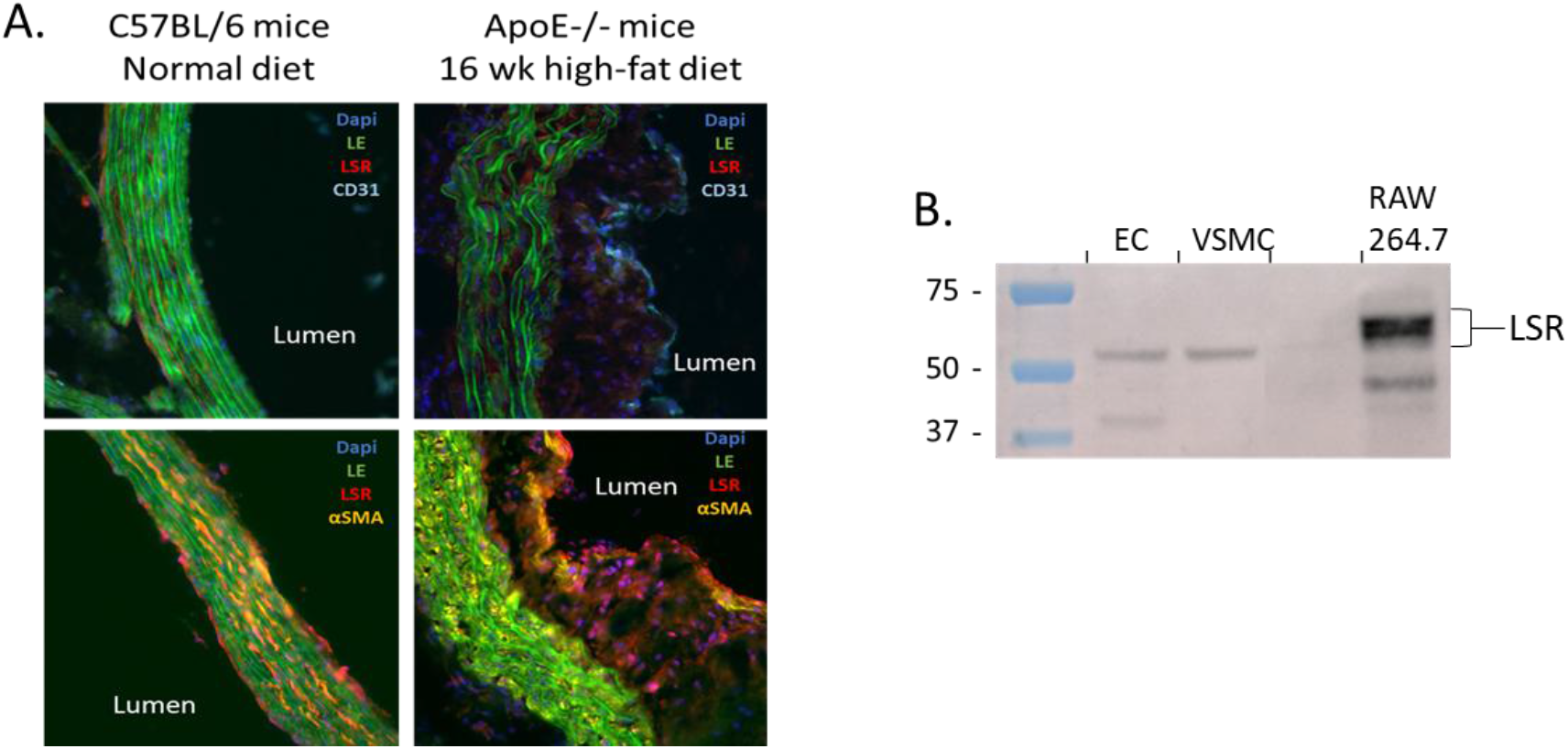
Detection of LSR in arterial vessel wall and atherosclerotic plaque. A) Immunofluorescence (IF) of LSR (red) in the aortas of C57BL/6 mice (left panels) on a normal diet and ApoE-/- mice after 16 weeks on a high-fat diet (right panels). Staining for CD31 (blue) in endothelial cells and α-SMA (yellow) in smooth muscle cells is shown (EL, elastic lamina). B) LSR immunoblot was performed on lysates from murine cells including aortic endothelial cells (EC), vascular smooth muscle cells (VSMC), and RAW264.7 macrophage cell line.

Detection of LSR in the arterial wall was confirmed by Western blot analysis which showed detectable levels of LSR in lysates prepared from primary cultures of murine aortic VSMC and EC (**Fig 1B**). In addition, all three subunits (α, α’, and β) of LSR were found in non-polarized (M0) Raw 264.7 macrophage cell line (Fig 1B).

Activation with LPS and IFNγ led to increased expression of M1 markers iNOS (4-fold, p < 0.01) and MCP-1 (2-fold, p < 0.05), while activation with IL-4 and IL-13 led to increased expression of M2 markers CD206 (6-fold, p < 0.01) and Arg-1 (175-fold, p < 0.01)(**Fig. 2A**). RT-qPCR and immunoblot analyses revealed that both LSR mRNA (**Fig. 2B**) and protein (**Fig. 2C**) levels were reduced in M1 macrophages (iNOS+ and MCP1+) compared to M2 macrophages (CD206+ and Arg-1+).

**Figure 2.**
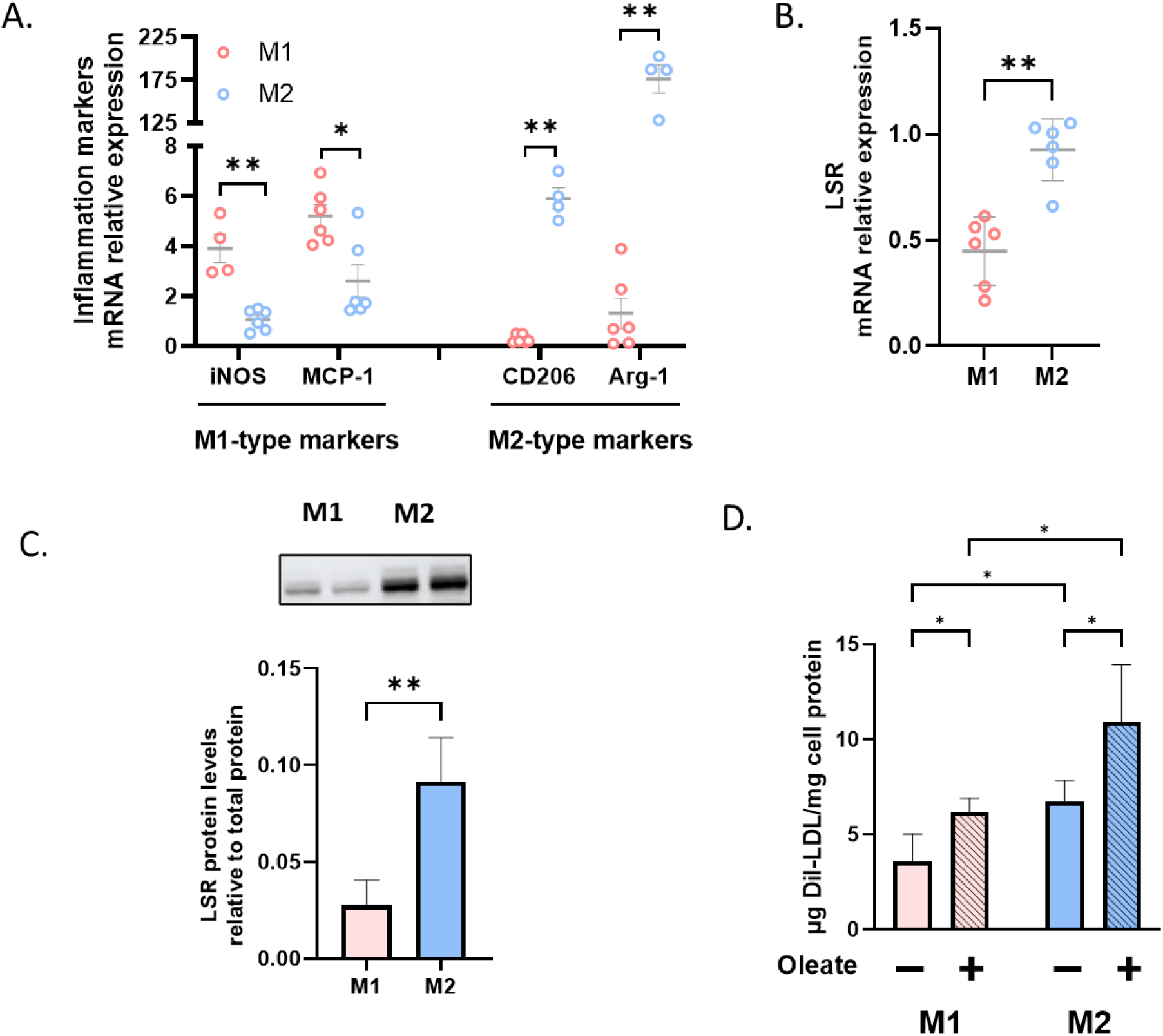
Effect of polarization of Raw 264.7 on inflammation markers (A), LSR mRNA (B) and LSR protein (C), as well as LSR activity. **(D)**. RAW264.7 macrophages were incubated 72 h at 37°C in the presence of LPS and IFNγ or IL4 and IL13 to induce pro-inflammatory (M1) or pro-resolving (M2) phenotypes. After which, cells were lysed for qPCR analysis of A) macrophage phenotype markers or B) LSR expression levels. Relative expression is shown as compared to M0 macrophages, and β-actin was used as the reference housekeeping gene C) Immunoblots were performed to detect LSR protein in cell lysates from M1 and M2 macrophages. D) To measure LSR activity, polarized cells were incubated 2h at 37°C in the absence or presence of 0.1 mM oleate with DII-LDL; fluorescence was measured in cell lysates. Specific DII-LDL internalization was calculated by subtracting nonspecific internalization measured in presence of 20-fold excess LDL. Results are shown as mean ± SD (N= 3, n = 2-3; Student’s t-test or one-way Anova plus Sidak’s multiple comparison test, * p < 0.05, ** p<0.01, **** p<0.0001).

We next sought to measure LSR activity first in M0 macrophages. Cells were incubated 2h at 37°C with increasing concentrations of DII LDL. Analysis revealed significantly higher amounts of specific oleate-induced internalized DiI-LDL which represents LSR activity after incubation with 50 µg/mL DiI LDL (**Supplementary Figure 1**). The same assay was performed after polarization of macrophages and revealed LSR activity as oleate-induced DiI LDL uptake in both M1 and M2 cell phenotypes. However, in both the absence or presence of oleate, the amount of DiI-LDL was significantly lower in the M1 cells as compared to M2 cells (**Fig. 2D**). Thus, these results showed that LDL uptake was significantly lower in M1 cells, and that the expression, protein levels, and activity of LSR as a lipoprotein receptor were reduced in pro-inflammatory M1 macrophages as compared to pro-resolving M2 macrophages.

To determine the consequences on macrophage lipid status in absence of LSR, siRNA was used to suppress *lsr* expression. *Lsr* knockdown by siRNA in Raw 264.7 M0 cells led to decreased expression of genes involved in lipoprotein uptake including LDL-R and the LDL-R related protein (LRP1) and transport (ABCA1)(**Fig. 3**). However, *lsr* knockdown did not significantly modify expression of M1 or M2 inflammatory markers (**Fig 3**).

**Figure 3.**
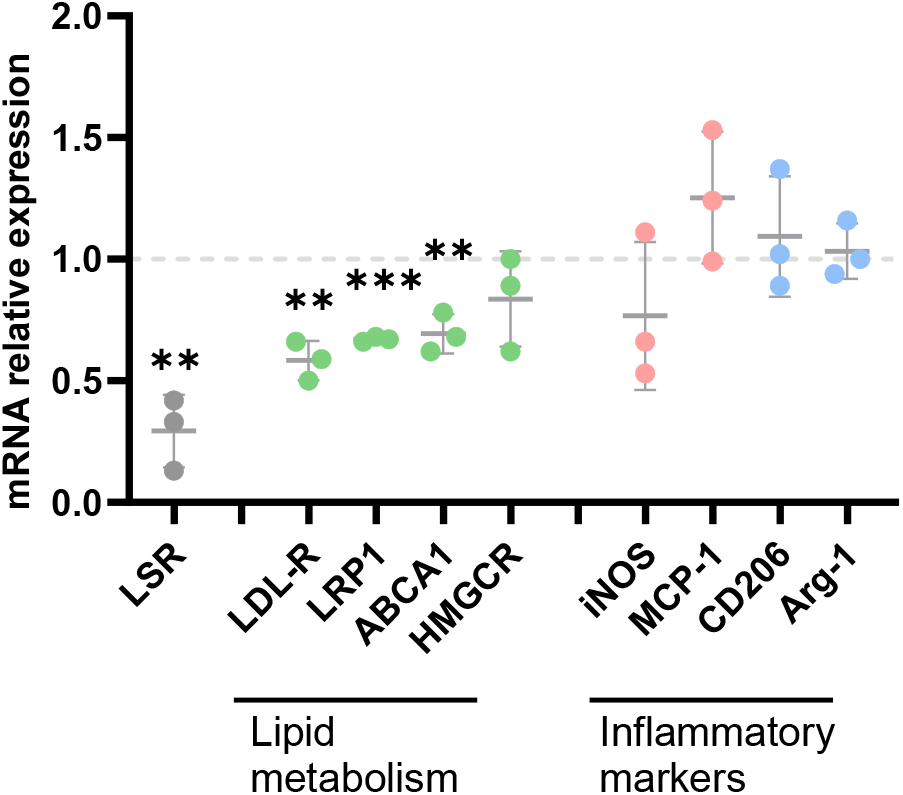
Effect of siRNA-mediated lsr knockdown in Raw 264.7 cells on relative expression lipid-related genes and inflammatory markers determined by RT-qPCR (β-actin was used as the reference housekeeping gene). Results are shown as mean ± SD (N=3, n=3; for qPCR results, Student’s t-test compared to control-siRNA treated cells for each gene, ** p<0.01, *** p<0.001).

Despite the observed changes in expression of genes involved in lipid metabolism, analysis of cell lipid composition did not reveal any detectable significant changes in cell cholesterol or triglycerides (**Table 1**).

**Table 1.** Cellular lipid content in RAW264.7 after siRNA-mediated *lsr* knockdown.

| Cell Lipid content (µg/well) | Control siRNA | LSR siRNA |
| --- | --- | --- |
| Total cholesterol (TC) | 7.33 ± 1.36 | 6.13 ± 2.31 |
| Cholesteryl esters (CE) | 0.33 ± 0.57 | 0.14 ± 0.25 |
| Free cholesterol (FC) | 7.76 ± 0.38 | 6.64 ± 1.33 |
| Triglycerides (TG) | 91.48 ± 7.37 | 109.05 ± 25.55 |
| Phospholipids (PL) | 10.21 ± 0.46 | 8.79 ± 1.22 |
| Lipid ratio |  |  |
| TC/PL | 0.72 ± 0.15 | 0.68 ± 0.18 |
| CE/PL | 0.03 ± 0.06 | 0.01 ± 0.02 |
| FC/PL | 0.76 ± 0.02 | 0.75 ± 0.05 |
| TG/PL | 8.98 ± 0.34 | 12.99 ± 5.15 |

Metabolic activity was next measured using the Seahorse in a parallel set of cells following siRNA-mediated *lsr* knockdown (**Fig. 4**); the level of knockdown was verified (84 ± 5%, n = 2). Measurement of glycolysis rate (**Fig. 4A**) revealed that *lsr* knockdown led to a significant increase in both basal glycolysis (**Fig. 4B**), as well as increased % proton efflux rate (PER; **Fig. 4C**). This contributed to a significantly lower ratio between mitochondria-derived oxygen consumption rate and glycolysis-derived PER (**Fig. 4D**). The increased glycolytic rate was confirmed using the glycolysis stress test (**Fig. 4E,F**), which involves a first step of starvation for 20 min, followed by addition of glucose. There was a tendency towards increased glycolytic capacity (**Fig. 4G**), which was measured by the addition of oligomycin to inhibit oxidative phosphorylation, but this did not reach statistical significance. However, glycolytic reserve or the capacity of the cell to respond to energetic demand was significantly reduced (**Fig. 4H**). Together, these results showed significant metabolic shift towards glucose-based metabolic changes following siRNA-mediated *lsr* knockdown in macrophages.

**Figure 4.**
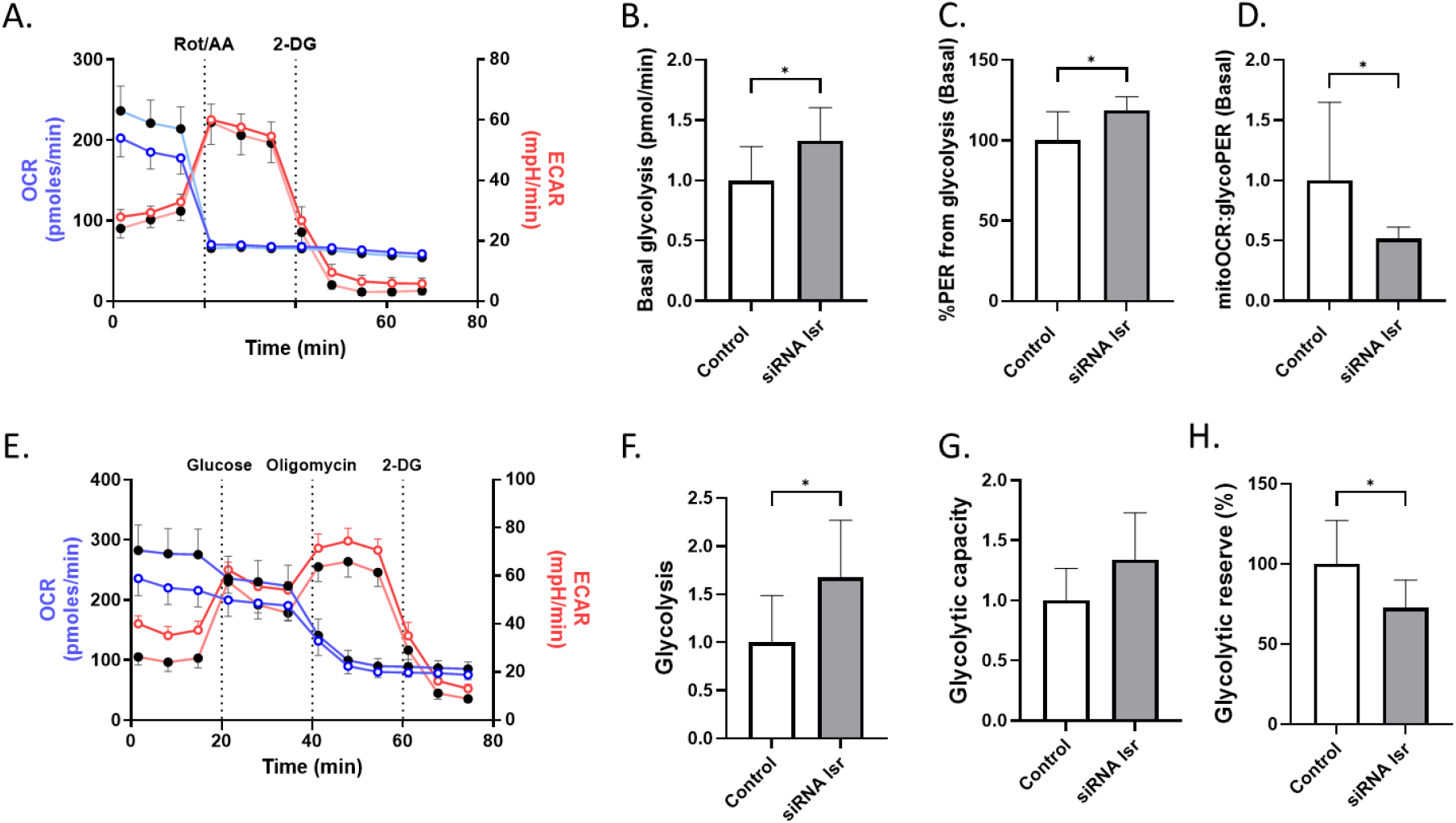
Effect of siRNA-mediated lsr knockdown on cell metabolic state. RAW264.7 cells were treated with control (open circles) or lsr (closed circles) siRNA as described in Fig. 3. Glycolysis rate was measured using the Seahorse glycolytic rate (A-D) and glycolytic stress assays (E-H). Results are shown as mean ± SD (N=3, n=3; * p < 0.05, Student’s t-test as compared to control). ECAR, extracellular acidification rate; OCR, oxygen consumption rate; PER, proton efflux rate; 2-DG, 2-deoxyglucose.

siRNA-mediated *lsr* knockdown did not lead to a significant change in glycolysis-derived ATP production, but rather increased rate of ATP production derived from oxidative phosphorylation (**Fig. 5A-C**). This led to a significantly increased ATP rate index, which represents the ratio of ATP production derived from oxidative phosphorylation to that derived from glycolysis. Total ATP production rate difference due to *lsr* knockdown did not achieve statistical significance.

**Figure 5.**
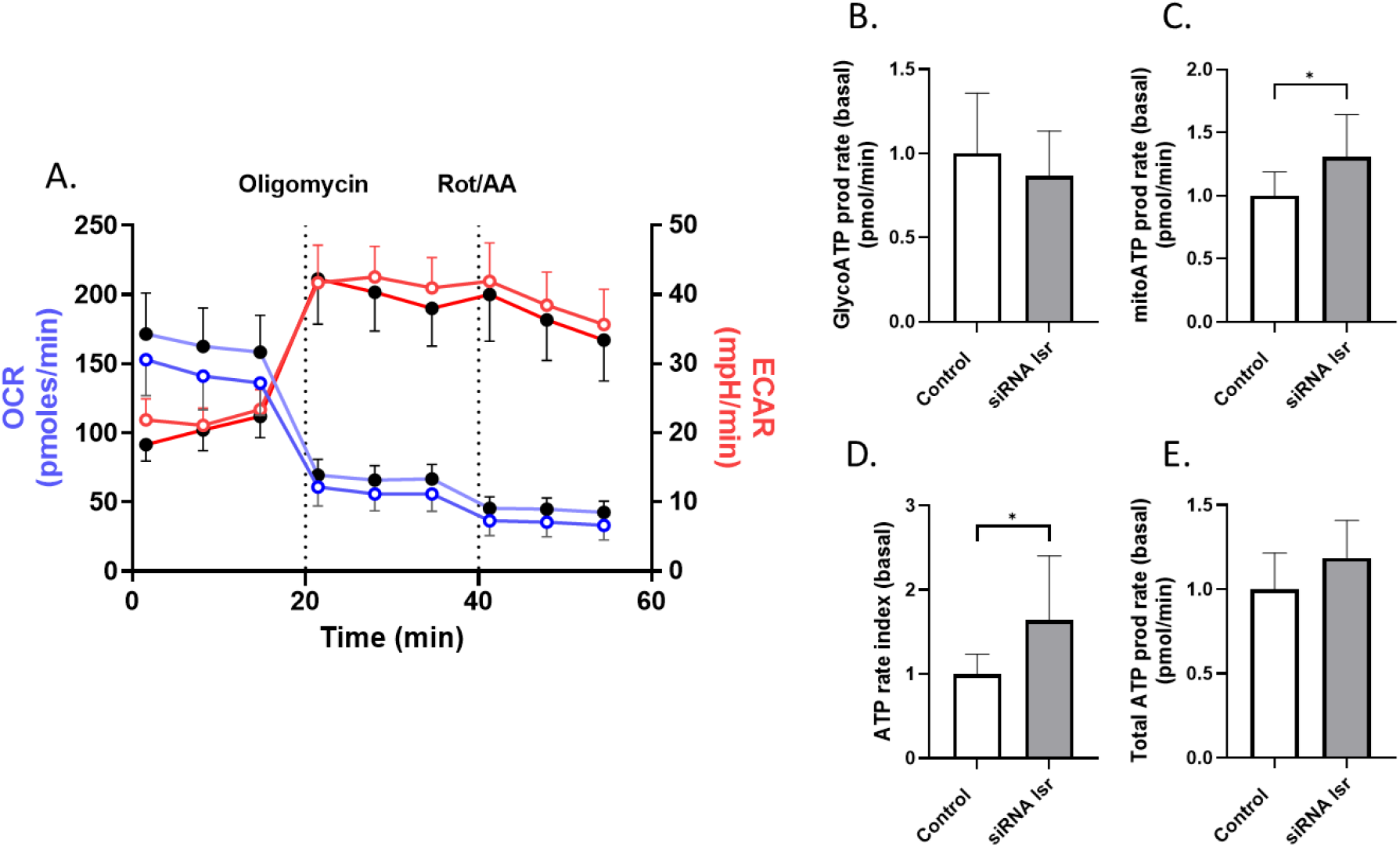
Effect of siRNA-mediated lsr knockdown on cell ATP production. RAW264.7 cells were treated with siRNA-control (open circles) or -lsr (closed circles) as described in Fig 3. ATP production was measured using the Seahorse® assay. Results are shown as mean ± SD (N=3, n=3; * p < 0.05, Student’s t-test as compared to control). ECAR, extracellular acidification rate; OCR, oxygen consumption rate).

Together, these results indicated significant changes in metabolic programming following siRNA-mediated *lsr* knockdown.

### LSR activity in foam cells

To determine if LSR as lipoprotein receptor could participate in the uptake of oxidized LDL, thus contributing to foam cell formation, DiI-LDL was minimally oxidized by UV (**Fig. 6A**), and then incubated (50 µg/mL) for 2h at 37°C with RAW264.7 macrophages to assess lipoprotein uptake capacity in the absence or presence of oleate. Results showed that while oleate-induced DiI-LDL uptake was increased as previously observed (**Fig. 6B**), the uptake of DiI-oxLDL in the presence of oleate was much lower as compared to what was observed for DiI-LDL (**Fig. 6C**), suggesting that affinity of oxLDL for LSR is significantly reduced as compared to LDL.

**Figure 6.**
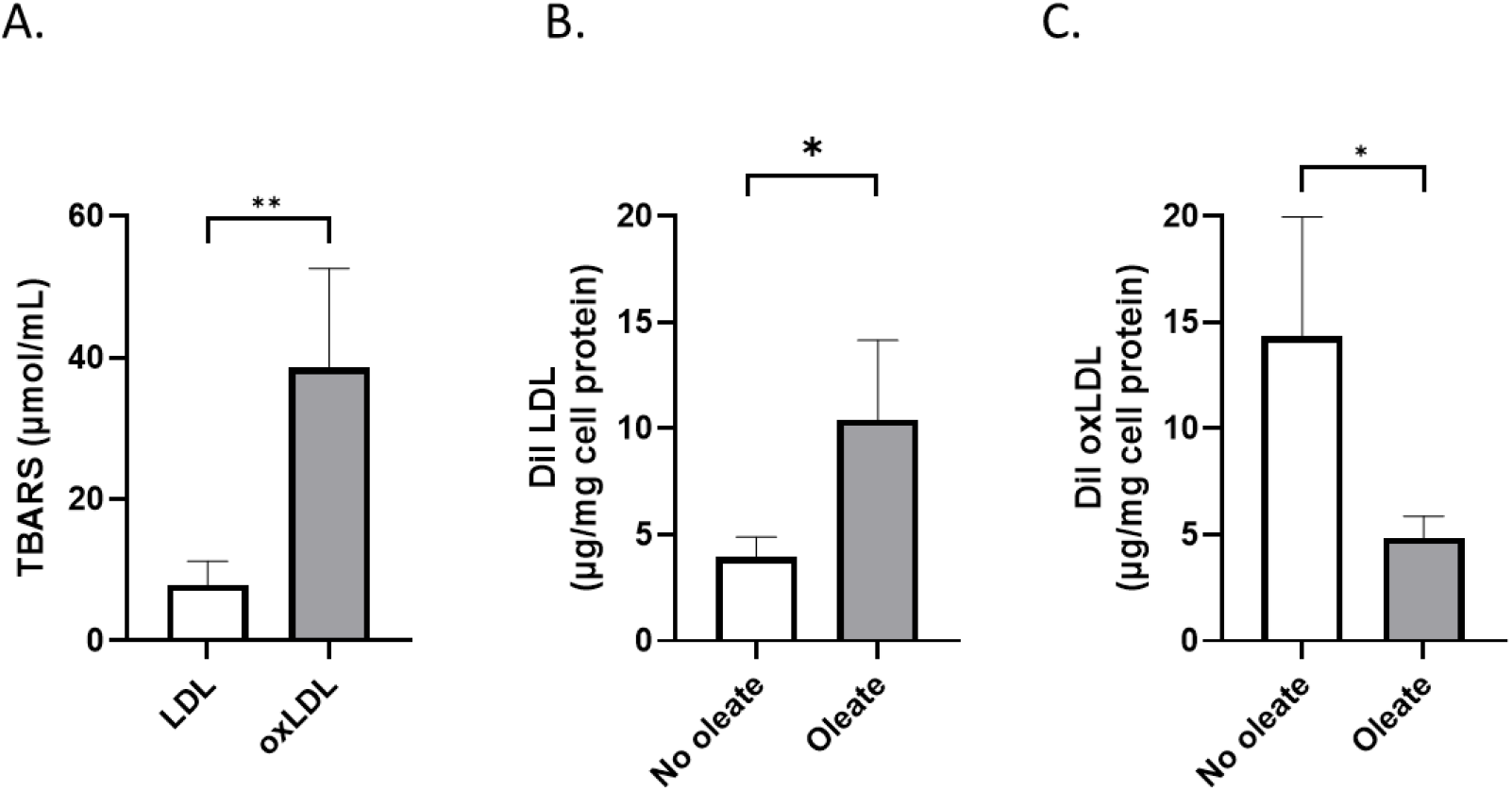
Uptake of oxidized LDL (oxLDL) by LSR. DiI-LDL was minimally oxidized by UV as described in Materials and Methods, as determined by TBAR assay (A). Amount of lipoprotein internalized in RAW 264.7 macrophages after incubation 2h at 37°C with 50 µg/mL DiI LDL (B) or DiI oxLDL (C) in absence or presence of 0,1 mM oleate. Results are shown as mean ± SD (N = 3 independent experiments, n = 3)(* p < 0,05, ** p < 0,01, using Student’s t-test).

To determine the effect of foam cell formation on macrophage lipid handling, cells were incubated 24h at 37°C with 100 µg/mL LDL or oxLDL in LPDS-containing medium. Cells maintained in growth medium (complete) were used as controls. LSR mRNA levels were significantly down-regulated following exposure of macrophages to oxLDL as compared to either control cells or those in LPDS (**Fig. 7A**), while those of LDL-R were significantly lower, but only when compared to cells in LPDS (**Fig. 7B**). Unlike LSR, LDL expression was also down-regulated following exposure of cells to LDL, as compared to macrophages incubated in complete or LPDS medium. Measurement of LSR activity showed significantly lower amounts of DiI-LDL internalized in the presence of oleate, but only when compared to control cells in complete medium (**Fig. 7C**). Cells in LPDS medium also displayed significantly lower LSR activity as compared to control cells. In absence of oleate, DiI-LDL uptake was reduced in all conditions as compared to control cells (**Fig. 7D**). These results indicate that pretreatment of cells with oxLDL significantly suppressed both LSR and LDL-R expression levels and macrophage capacity for lipoprotein internalization either in the absence or presence of oleate.

**Figure 7.**
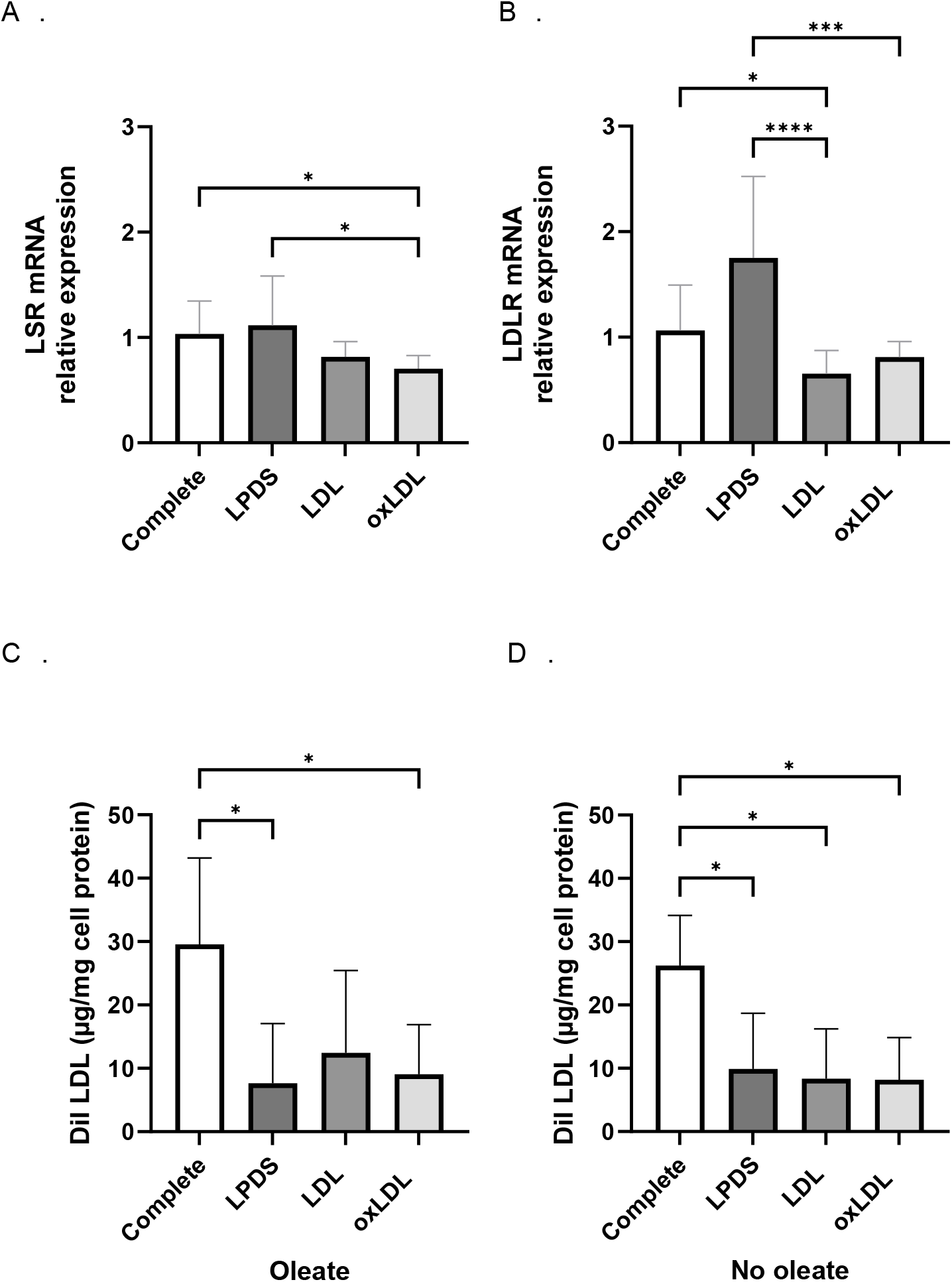
Effect of 24h pretreatment of RAW264.7 macrophages with oxLDL on DiI-LDL internalization in the absence or presence of oleate. Cells were incubated 4h at 37°C with medium containing 0.2% BSA, and then 24h at 37°C in the absence or presence of 100 µg/mL LDL or ox LDL in lipoprotein deficient serum-containing medium (LPDS) as described in Materials and Methods. Cells incubated in normal growth medium containing fetal bovine serum (Complete) were used as controls. One set of cells was used to determine LSR (A) and LDLR (B) mRNA levels. In another set of cells, DiI-LDL uptake was measured in the presence (C) or absence (D) of 0,1 mM oleate as described in Fig 2. Results are shown as mean ± SD (N = 3 with triplicates for qPCR and N = 4 independent experiments for DiI-LDL assay; * p < 0,05, *** p < 0,001, **** p < 0,0001, using one-way ANOVA, with comparison as indicated).

## Discussion

The results of this study demonstrate that polarization of Raw 264.7 macrophages to the pro-inflammatory M1 phenotype led to down-regulation of LSR mRNA and protein levels. DiI-labeled LDL uptake in the presence of oleate is a measure of LSR activity of lipoprotein receptor [7,10,17] and was also found to be significantly diminished in M1 macrophages as compared to M2 cells. LSR is unique in that it is the only known lipoprotein receptor to bind and endocytose LDL in the presence of oleate; oleate-induced LDL uptake was shown to be both LDL-R and LRP1-independent [17]. In addition, the LDL-R is unable to bind LDL in the presence of fatty acids [18]. The presence of fatty acids leads to conformational change of the LSR multimeric protein complex, exposing a binding site for ApoB or ApoE of the lipoprotein [19]. Upon binding of lipoproteins to LSR, this receptor mediates endocytosis of lipoproteins leading to lysosomal degradation of the particle. The presence of all 3 subunits α, α’ and β of LSR were detected in this cell line and it was previously demonstrated that at least 2 subunits are required for lysosomal degradation following LSR-mediated endocytosis [8].

We have shown that LSR displays higher affinity for triglyceride-rich lipoproteins including VLDL and chylomicron remnants, but is also able to bind LDL [7,17]. Because LDL contains only one apolipoprotein, apoB, and is less heterogeneous than VLDL and chylomicrons, it represents a useful ligand for testing receptor activity. Here, a density range of 1.025 < d < 1.055 to prepare LDL was used to avoid any potential contamination by VLDL/IDL or HDL. We previously demonstrated LSR mediated oleate-induced uptake of ^125^I-LDL in fibroblasts and hepatocytes as low as 0.2 mM. [17], and that this measurement was LSR-specific demonstrated by siRNA-mediated silencing of LSR [10]. Here, concentrations of 0.1 mM oleate used were non-toxic for the cells, as demonstrated by lack of cell protein loss (data not shown), and was sufficient for maximal inhibition of LDL binding to the LDL-R [18]. Therefore, we conclude that inflammatory stimuli of macrophages into M1-phenotype leads to a significant downregulation of LSR expression and protein levels, which translates into decrease lipoprotein uptake via the LSR pathway.

In the initial phase of atherosclerosis, lipoproteins accumulate in the sub-endothelial space, and are taken up by macrophages either as native or modified form via lipoprotein or scavenger receptors (CD36 or SRB1), respectively. Our data revealed lower oleate-induced uptake of DiI-oxLDL as compared to that when oleate was absent, which would suggest lower affinity of LSR for oxLDL as compared to native unmodified LDL. Another potential explanation is that scavenger receptor activity was significantly lower in the presence of oleate, at least under these assay conditions. The scavenger receptor CD36 has been shown to be involved in transport of long chain fatty acids including oleate [20], and that CD36 binding of oxLDL is fatty acid-dependent [21]. Further ligand-receptor interaction studies to determine relative affinities for oleate and oxLDL will help to clarify the role of each receptor in oxLDL uptake in the presence of oleate.

siRNA-mediated *lsr* knockdown led to significant consequences in the macrophages with regards to lipid metabolism. LDL-R and LRP1 as well as ABCA1 were downregulated, which would suggest a common regulatory step following *lsr* knockdown that leads to a general shutdown of cell lipoprotein trafficking. Decreased LSR and LDL-R would translate into lower ability to internalize lipoproteins and thus decreased lipid influx, while lower lipid output would result from decreased ABCA1-mediated secreted transport of lipoproteins into the extracellular medium. However, these modifications of genes involved in lipoprotein uptake and transport did not lead to any detectable significant changes in cell lipid levels after a 3-day siRNA treatment. It is possible that longer incubation times may be required to observe any modifications since significant changes in cell cholesterol levels in astroglial cells were observed following LSR silencing, but after a longer period of time (5 days)[14]. Interestingly, no significant changes were observed in expression of the M1 markers iNOS and MCP-1, or the M2 markers CD206 and Arg1, demonstrating that this LSR-mediated change in lipid metabolism genes occurred without polarization of macrophages into a specific phenotype.

On the other hand, analyses of metabolic activity using Seahorse® assay revealed significant metabolic changes following *lsr* knockdown. Glycolytic activity was increased, confirmed by increased PER which is derived from glycolysis, and is correlated with extracellular lactate levels. This contributed to a lower ratio of mitochondria-derived oxygen consumption rate and glycolysis-derived PER. Placing cells on a glycolysis stress test also confirmed increased glycolysis, which most likely caused the decrease in glycolytic reserve and thus its capacity to respond to energy demands. Despite this change in glycolysis, there was not a significant change in glycolysis-derived ATP production. Instead, there was an increased ATP production rate derived from oxidative phosphorylation, which may have resulted from increased glycolysis-derived pyruvate flux that enters the Kreb’s cycle. This did not, however, result in a significant change in total ATP production rate. Therefore, although no significant changes in cell lipid composition were observed, the downregulation of LDL-R, LRP1 and ABCA1 could be considered as part of a general shutdown in lipoprotein uptake and transport, thereby promoting the metabolic shift to glycolysis rather than lipid-derived energy production. Since glycolysis is the metabolic state of pro-inflammatory macrophages, the lipid dyshomeostasis induced by reduced LSR may render the cells more susceptible to pro-inflammatory stimuli. This remains to be confirmed.

Foam cell formation induced by 24h incubation with oxLDL also led to both a significant down-regulation of LSR mRNA levels, as well as decreased LSR activity demonstrated by lower oleate-induced DII-LDL uptake. It was previously reported that ox-LDL-induced foam cell formation leads to increased uptake of F-fluorodeoxyglucose and increased glycolysis in macrophages [21], that could be attributed in part to ox-LDL/CD36 signaling [22]. Here, our results show that LSR activity is reduced in foam cell formation, which could contribute to a metabolic shift towards glycolysis, as suggested by the increased glycolysis observed following siRNA-mediated LSR silencing.

To summarize, these studies demonstrated that LSR expression and activity were related to macrophage metabolic activity and phenotype. LSR deletion resulted in inhibition of lipid metabolism genes, as well as an increase in glycolytic activity. Therefore, one would expect that the absence of LSR would make macrophages more likely to evolve to the pro-inflammatory M1 phenotype. In addition, reduced lipid utilization could promote a focus on lipid storage, thereby leading to lipid-loaded cells, which is a key step in the early stages of atherosclerotic plaque formation. Our data also showed significantly lower levels of internalized DiI-LDL in absence of oleate in M1 macrophages and after oxLDL-induced foam cell formation. This would suggest that both LSR and LDLR expression and activities are closely linked to the metabolic state of the cell. Pro-inflammatory stimuli, or oxLDL would thus induce a coordinated response which includes a general shutdown of native lipoprotein uptake and transport, as well as a metabolic shift from fatty acid oxidation to glycolysis. However, neither of these 2 factors are necessarily needed since siRNA-mediated knockdown of the LSR alone can signal the macrophage to shift to glycolysis. Further investigation is required to determine the signaling pathway that links LSR to the regulation of metabolic programming in macrophages. These results provide reveal an important link between LSR, macrophage lipid homeostasis and metabolic programming, which may provide new insight on understanding how immune cells dynamically adapt their function in response to tissue microenvironments. Indeed, lipoprotein receptors can be viewed as gateway hubs that regulate lipid influx thereby influencing macrophage fate, whether it is driving the cells towards an M1 or M2 phenotype. Targeting these pathways may provide new therapeutic avenues for controlling macrophage polarization thereby reducing chronic inflammatory processes in atherosclerosis.

## Acknowledgements

The authors thank Dr Véronique Regnault for her expert advice in interpretation of the Seahorse® data, and for kindly providing the mouse VSMC and EC. GV is recipient of a French Ministry of Health, Education and Space MESRE thesis fellowship. Funds for this project were provided in part by the MESRE. Supported by CARE-IN-HEALTH funded by the European Union within HORIZON EUROPE Health Framework Program under grant agreement 101 095 413. Views and opinions expressed are however those of the authors and do not necessarily reflect those of the European Union or the European Health and Digital Executive Agency. and the Horizon European project Care-in-Health.

