## Supplementary data for "Role of the lipolysis stimulated lipoprotein receptor LSR in the regulation of macrophage lipid homeostasis and metabolic activity"

### Methods

**Supplementary Table 1**. Antibodies used

| Antibody | Supplier | Reference | Dilution* |
|---|---|---|---|
| Anti-LSR | US Biological Life Sciences | L25052057 | 1:200 (IF) |
| Anti-CD68 | Abcam | AB53444 | 1:200 (IF) |
| Anti-MCP-1 | Cell Signaling | 2027S | 1:200 (IF) |
| Anti-CD206 | Biorbyt | Orb308745 | 1:200 (IF) |
| Anti-CD31 | Abbiotec | 250590 | 1:200 (IF) |
| Anti-αSMA | Merck Sigma | A5228 | 1:200 (IF) |
| Anti-rabbit IgG (555) | Invitrogen | A21429 | 1:200 (IF) |
| Anti-rabbit IgG (488) | Invitrogen | A21206 | 1:200 (IF) |
| Anti-rat IgG (555) | Invitrogen | A21094 | 1:200 (IF) |
| Anti-rat IgG (488) | Invitrogen | A21094 | 1:200 (IF) |
| Anti-mouse IgG (488) | Invitrogen | A21200 | 1:200 (IF) |
| Anti-mouse IgG (555) | Invitrogen | A21127 | 1:200 (IF) |
| HRP-conjugated Anti-rabbit IgG | GE Healthcare | NA931V | 1:1000 (WB) |
| HRP-conjugated Anti-mouse IgG | GE Healthcare | NA934V | 1:1000 (WB) |

*IF, immunofluorescence; WB, Western blot

**Supplementary Table 2.** qPCR Primers

| Gene | Forward Primer (5’-3’) | Reverse Primer (5’-3’) |
|---|---|---|
| *lsr* | AGTAATACACTCCACTGTCTCCCCAG | CAGGAGAATCACCATCACAGGAA |
| *ldl-r* | TGGCTATACCTACCCCTCAAGACAG | GATCCCGGAAAGAGACGGAT |
| *lrp-1* | CGAGAGCCTTTGTGCTGGATGA | CGGATGTCCTCAATGAGGG |
| *abca1* | CAACCCCTGCTTCCGTTATCCAA | GAGAACAGGCGAGACACGATGGAC |
| *hmgcr* | CCCCACATTCACTCTTGACGCTCT | GCTGGCGGACGCCTGACAT |
| *inos* | GACATTACGACCCCTCCCAC | GCACATGCAAGGAAGGGAAC |
| *mcp-1* | GCTACAAGAGGATCACCAGCAG | GTCTGGACCCATTCCTTCTTGG |
| *cd206* | CTAACTGGGGTGCTGACGAG | GGCAGTTGAGGAGGTTCAGT |
| *arg-1* | AATGAAGAGCTGGCTGGTGT | CTGGTTGTCAGGGGAGTGTT |
| *β-actin* | GCAGGAGTACGATGAGTCCG | ACGCAGCTCAGTAACAGTCC |

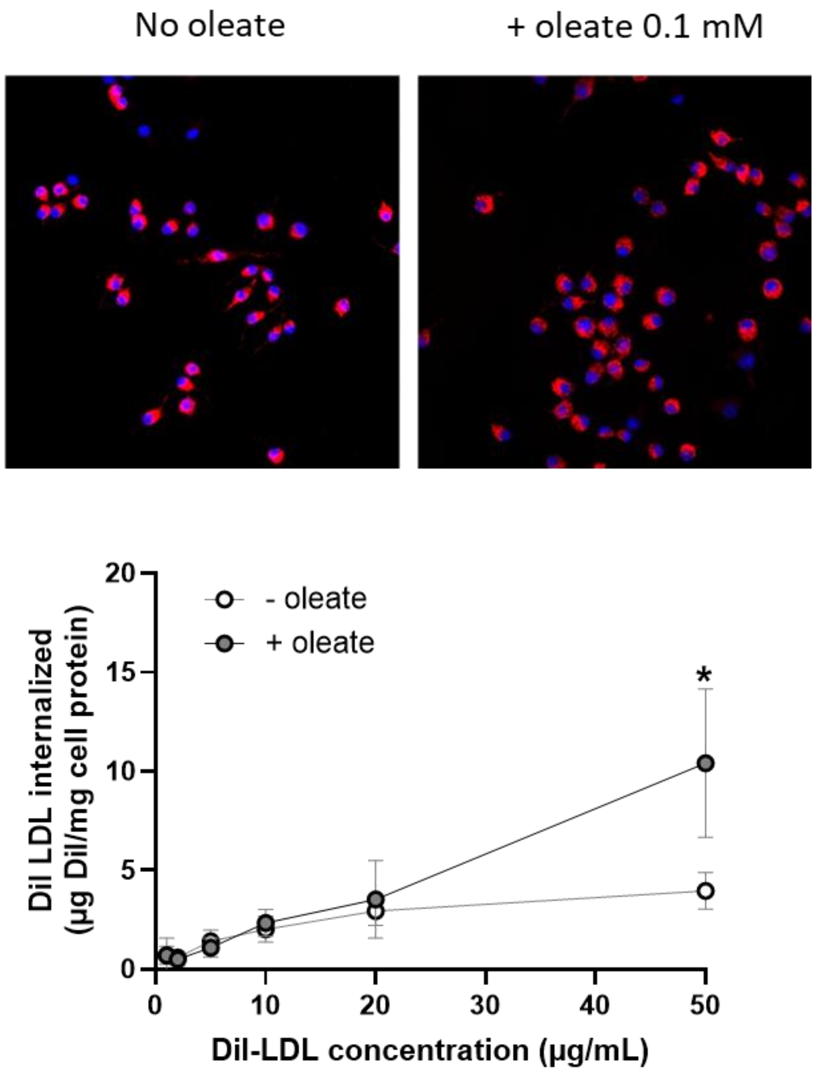


**Supplementary Figure 1.** LSR activity in non-polarized RAW264.7 macrophages. Cells (80% confluence) were incubated 2h at 37°C with DiI-LDL at the indicated concentrations in the absence and presence of 0.1 mM oleate. Non-specific binding was measured in one set of cells incubated with 20-fold excess unlabeled LDL to measure non-specific binding and subtracted from the total fluorescence to obtain specific amount of internalization, indicated as DII LDL internalized (µg DiI-LDL/mg cell protein) (N = 3, n =3; * $p < 0.05$ t-test). Representative images are shown of cells incubated 2h at 37°C with 10 µg/mL DiI-LDL.
